# Local nucleation of microtubule bundles through tubulin concentration into a condensed tau phase

**DOI:** 10.1101/119800

**Authors:** Amayra Hernández-Vega, Marcus Braun, Lara Scharrel, Marcus Jahnel, Susanne Wegmann, Bradley T. Hyman, Simon Alberti, Stefan Diez, Anthony A. Hyman

**Affiliations:** Max Planck Institute of Molecular Cell Biology and Genetics, Dresden, Germany; B CUBE – Center for Molecular Bioengineering, Technische Universität Dresden, Dresden, Germany; BIOTEC, Technische Universität Dresden, Germany; Dept. Neurology, Massachusetts General Hospital, Harvard Medical School, Boston, MA

## Abstract

Non-centrosomal microtubule bundles play important roles in cellular organization and function. Although many diverse proteins are known that can bundle microtubules, biochemical mechanisms by which cells could locally control the nucleation and formation of microtubule bundles are understudied. Here, we demonstrate that concentration of tubulin into a condensed, liquid-like compartment composed of the unstructured neuronal protein tau is sufficient to nucleate microtubule bundles. We show that under conditions of macro-molecular crowding, tau forms liquid drops. Tubulin partitions into these drops, efficiently increasing tubulin concentration and driving the nucleation of microtubules. These growing microtubules form bundles enclosed in a liquid sheath of tau. Our data suggest that condensed compartments of microtubule bundling proteins could promote the local formation of microtubule bundles in neurons by acting as non-centrosomal microtubule nucleation centers, and that liquid-like tau encapsulation could provide both stability and plasticity to long axonal microtubule bundles.

## INTRODUCTION

In order to organize their microtubule arrays, neurons must solve a number of challenges. First, they must drive the nucleation of microtubules in a centrosome independent manner. Secondly, they must organize microtubules into bundles, which can be millimeters long. Although there is considerable knowledge about how neurons use microtubule bundles for microtubule-based transport, we have less knowledge of how microtubule bundles form in the first place (Baas et al., 2005; Chen et al., 2017; Sanchez and Feldman, 2016; Tanaka and Kirschner, 1991). More generally, we have little information on how cells locally nucleate microtubules in a centrosome independent manner (Matamoros and Baas, 2016; Sánchez-Huertas et al., 2016).

The local formation of microtubule asters has been well studied and is controlled by nucleation of microtubules at centrosomes. Here, it is thought that concentration of proteins at centrosomes can drive local nucleation of microtubule asters. Indeed, recent work suggests that tubulin concentration at centrosomes could be a key mechanism driving aster nucleation (Woodruff et al., 2017, under review, bioRxiv preprint). However, the local nucleation of microtubule bundles is much less studied, and we know little about the cell biology of this problem. What sorts of biochemical mechanisms could drive local nucleation and formation of microtubule bundles?. The neurons are rich in unstructured microtubule-associated proteins. An important component of the microtubule cytoskeleton in neurons is the protein tau. Tau, also known as MAPT (Microtubule Associated Protein Tau), is found aggregated in a number of neurodegenerative disorders including Alzheimer’s disease and has been also genetically linked to some of these diseases (Arendt et al., 2016). In healthy brains, tau localizes to the axons of neurons while other MAPs localize to dendrites (Aronov et al., 2001; Matus, 1988). *In vitro*, it can stimulate the growth rate of microtubules (Drechsel et al., 1992; Kadavath et al., 2015), diffuse along and bundle microtubules (Chen et al., 1992; Chung et al., 2016; Hinrichs et al., 2012; Rosenberg et al., 2008).

Tau consists of up to four microtubule binding domains surrounded by intrinsically disordered regions (Fig. 1A and Mukrasch et al., 2009). Recent work suggests that many intrinsically disordered proteins can form liquid drops by demixing from buffer (Brangwynne et al., 2009; Elbaum-Garfinkle et al., 2015; Pak et al., 2016; Patel et al., 2015). Once these drops are formed, other client proteins can partition into the drops, where they are concentrated, with the degree of concentration depending on the partition coefficient. This concentration of client proteins provides a potential mechanism to increase the rates of reactions (Banjade and Rosen, 2014; Jiang et al., 2015; Li et al., 2012; Su et al., 2016).

Here, we demonstrate that under conditions of molecular crowding, tau forms liquid drops. Tubulin partitions into these drops, where it nucleates and drives the formation of microtubule bundles. These bundles deform the drops, yet remain surrounded by a liquid sheath of tau. We suggest that tau drops can drive the formation of microtubule bundles in neurons by acting as non-centrosomal microtubule nucleation centers and that tau liquid-like encapsulation could provide both stability and plasticity to long axonal microtubule bundles.

## RESULTS

We wanted to establish a system in which a neuronal microtubule-associated protein forms liquid drops by phase separation. To investigate whether tau can form drops, we first expressed the protein in insect cells and purified it by affinity chromatography (Fig. S1A). A solution of tau is diffuse and forms no obvious structures. When we added molecular crowder, tau phase-separated to form drops (Fig. 1B and Fig. S1B). We confirmed the liquid-like nature of these drops by showing (i) they rapidly fused and relaxed into bigger drops (Fig. 1C and movie 1), (ii) they rapidly recovered their fluorescence after photo-bleaching, indicating that the protein in the drops is highly diffusible (Fig. 1D and movie 2), (iii) they wetted onto glass surfaces (Fig. 1E and movie 3), and (iv) they underwent fission (movie 3).

To look at the interaction of tubulin with tau drops, we added tubulin in the absence of GTP. Under these conditions, tubulin partitioned into the drops (Fig. 2A). The amount of tubulin partitioning into the drops increased with increasing tubulin concentration (Fig. S2A) such that the concentration inside the drop was consistently more than ten-fold higher than outside the drop (Fig. 2B). As an example, when tubulin was added at an overall concentration of 1 μΜ, the concentration in the drop was higher than 20 μΜ (Fig. S2B). Interestingly, tubulin itself stimulated the formation of tau drops, suggesting a positive feedback mechanism between tubulin partitioning and tau drop formation (Fig. S3).

To assess the potential role of tau drops as nucleators for the growth of microtubules, we repeated the experiment in the presence of GTP by adding 5 μΜ tubulin together with 1 mM GTP to preformed tau drops. At this concentration, tubulin alone was unable to nucleate microtubules (Fig. S4). Within a few minutes after addition of tubulin-GTP, the tau drops deformed in a rod-like shape as microtubule bundles grow inside the drops (Fig. 2C and movie 4 and 5). Drop deformation was in most cases bidirectional (see Fig. 2D for a typical example). At longer times, dense networks of tau-encapsulated microtubule bundles formed, and no individual drops remained (Fig. 2E). We note that bundle formation in drops occurred under conditions where the tubulin concentration in solution was as low as 0.75 μΜ (Fig. 2F), more than an order of magnitude below the critical concentration for microtubule spontaneous nucleation in the absence of tau drops (Drechsel et al., 1992; Voter and Erickson, 1984; Wieczorek et al., 2015). In fact, in the absence of tau, regardless of the presence of crowding agents, 1–5 μΜ tubulin did not support microtubule nucleation or polymerization (Fig. S4, movie 6 and Wieczorek et al., 2013).

To further investigate the nature of the tau-encapsulated microtubule bundles, we looked at the relative dynamics of tubulin and tau by photo-bleaching. When we photo-bleached the microtubule bundles, little recovery of the tubulin fluorescence was observed (Fig. 3A and movie 7). This supports the idea that the microtubules within the tau drops formed stable bundles. When we photo-bleached tau, it rapidly recovered (Fig. 3A and movie 7), demonstrating that tau molecules around the microtubule bundles are freely diffusible. This suggests that after formation, the stable microtubule bundles are embedded in a viscous fluid of tau. Further evidence supporting the idea of a liquid behavior of this tau encapsulation comes from the observation that the tau-encapsulated microtubule bundles fused into larger assemblies (Fig. 3B and movie 8) and that newly formed tau drops were able to fuse with pre-existing tau-encapsulated microtubule bundles (Fig. 3C).

To examine how tau bundles microtubules, we added tau to preassembled, stabilized microtubules. Both, in the presence and absence of crowding agents, binding saturated at tau densities consistent with a mono-layer of tau around the microtubules (Fig. S5A,B), suggesting that more than one microtubule was necessary to form a thick liquid coating of tau. Tau-coated microtubules interacted with each other by zippering together (Fig. S5C and movie 9). This is consistent with recent work showing that tau binds to microtubules through its microtubule binding domains, while the intrinsically disordered N-terminal domain acts as a brush, which keep microtubules at a constant distance in microtubule bundles (Chung et al., 2016; Marx et al., 2000; Méphon-Gaspard et al., 2016; Mukhopadhyay and Hoh, 2001; Mukhopadhyay et al., 2004).

When we added tau drops containing tubulin and GTP to preassembled, stabilized microtubules, we noticed that tau drops bound and directed microtubule growth alongside the preexisting microtubules (Fig. 3D and movie 10). These observations indicate that tau, both as a thin layer around individual microtubules or as an encapsulating liquid embedding surrounding bundles, mediates attractive microtubule-microtubule interactions.

To examine the role of tau drops in stabilizing microtubules and microtubule bundles, we looked for conditions that would interfere with the interaction of tau and tubulin. For this purpose, we used heparin, a negatively charged polymer, which has been shown to interfere with the tau-tubulin interaction (Goedert et al., 1996). When we added heparin to tau-encapsulated microtubule bundles, tau de-wetted from the bundles, reforming spherical drops (Fig. 4A,B and movie 11). At the same time, the microtubules unbundled (Fig. 4A,B and movie 11), and depolymerized (Fig. S6). Presumably, tau drops facilitate the stabilization of microtubule bundles, either by keeping the tubulin concentration in the surrounding compartment above the critical concentration and/or by stabilizing the microtubule lattice by its multiple binding sites.

## DISCUSSION

Taken together, our experimental results show that tau can form liquid drops. Tubulin partitions into the drops, raising its local concentration above the critical concentration for nucleation of microtubules. These nucleated microtubules form bundles and grow, and collectively deform the tau drops into a rod-like shape tau-encapsulated microtubule bundled structure (see model in Fig. 4C). Tau’s multiple tubulin dimers binding sites, binding at the interface between tubulin hetero-dimers, together with its conformational flexibility may contribute to the nucleation of microtubules, and the formation of microtubule bundles (Kadavath et al., 2015; Li et al., 2015; Melo et al., 2016). Currently, we do not know whether similar mechanisms operate in vivo. Tau is known to localize to microtubule bundles in vivo, suggesting a role in organizing microtubule bundles. Although the mouse knock out of tau shows little phenotype, the deletion results in upregulation of another MAP, MAP2, suggesting that redundancy may prevent the observation of phenotype (Ma et al., 2014). A manuscript under review looking at the expression of GFP-tau constructs in neurons suggests that they also form higher order assemblies *in vivo* (Wegmann et al., under review). Although our work has focused on Tau, our experiments suggest that in principle, any microtubule bundling protein that forms drops could also act as a local site of microtubule bundle formation.

Our experiments require molecular crowders to drive the formation of tau drops at physiological salt concentrations. Presumably, this is substituting for a tau-drop nucleating factor that exists *in vivo*, that is not yet identified. Our results suggest that tubulin itself is part of the nucleation mechanism. Recently tau drops formation has been reported to form in the absence of a molecular crowder using RNA as nucleator under conditions of low salt and low pH, a tau-RNA association in vivo was also observed in this study (Zhang et al., 2017, bioRxiv preprint). The fact that tubulin can itself stimulate the formation of tau drops suggests an interesting feedback mechanism, in which most free tubulin in the axon could end up in drops. We can imagine that in axons, drop formation is stimulated when microtubule bundles are required for instance during neuron differentiation or regeneration. Another possibility is that drops are always present *in vivo* but that tubulin can only partition into them upon the removal of some inhibition signal, or that tubulin is always present in drops in a form in which microtubule nucleation is inhibited, and that removal of this inhibition would then drive bundle formation. More work on the formation of microtubule bundles in axons will be required to distinguish these possibilities.

Recent work on other unstructured proteins such as FUS has suggested that a liquid-solid phase transition is involved in the onset of protein aggregation in the motor neuron disease ALS (Patel et al., 2015). The fact that tau can also form liquid drops suggests that potentially, such liquid-solid phase transitions could be involved in the onset of Alzheimer’s disease.

Our work supports the idea that the local concentration of tubulin, driven by phase separation of regulatory factors could be a general mechanism driving the formation of microtubule arrays. In *Xenopus* extracts, the protein BuGZ has been shown to form liquid drops, into which tubulin can partition and polymerize microtubules, suggesting a mechanism for spindle formation (Jiang et al., 2015). A similar mechanism for promoting local actin polymerization by its recruitment into phase-separated compartments was recently also observed (Banjade and Rosen, 2014; Li et al., 2012; Su et al., 2016). These experiments, together with our findings, suggest that concentration of cytoskeletal monomers into liquid drops of cytoskeletal regulatory proteins could be a general mechanism for the formation of local nucleation centers for cytoskeletal filaments. The types of arrays that arise from this concentration mechanism, be it bundles or asters, would depend on the type of microtubule associated proteins that are present in the drops.

**Fig. 1.**
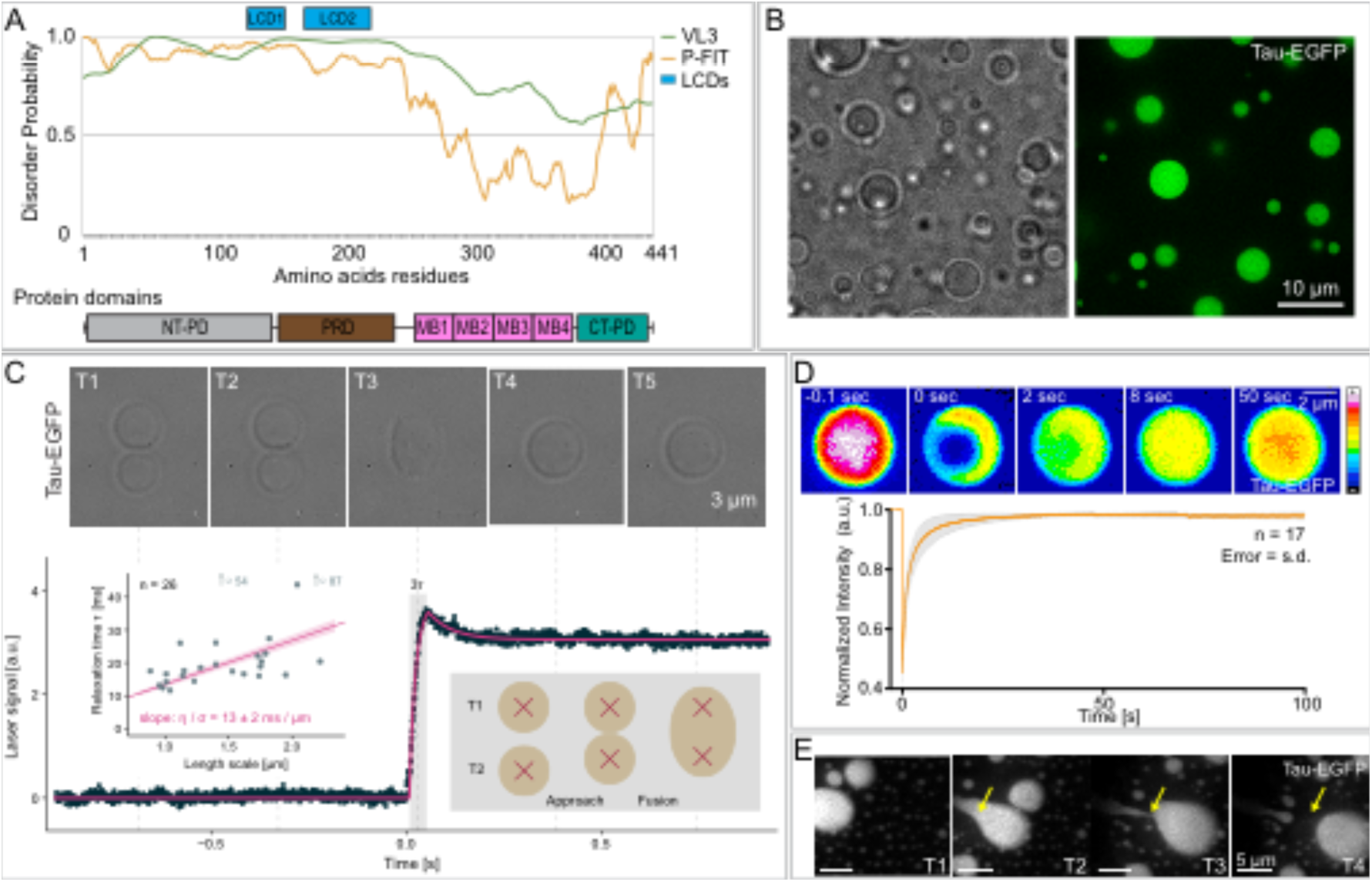
Tau phase separates into liquid-like drops. **(A)** Prediction of the degree of disorder along tau protein (htau441). PONDR-FIT (P-FIT) and VL3 algorithms are shown in orange and green respectively. A given protein region is considered to be disordered when the disorder probability is above a threshold of 0.5 (y axis). LCD1 and LCD2 (blue rectangles) highlight regions of the protein with potential low complexity. Protein domains along the protein are shown below being; NT-PD, N-terminal projection domain; PRD, proline-rich domain; MB1-4, microtubule binding repeats 1 to 4; CT-PD, C-terminal projection domain. **(B)** Tau drops formed *in vitro* in the presence of crowding agent. Brightfield and fluorescence microscopy images of tau-EGFP drops. See also figure S1. Tau drops were formed with 25 μΜ tau-EGFP, 25 mM Hepes, 150 mM KCl, 1 mM DTT, 10% dextran, pH 7.4, for all experiments in this figure. **(C)** Fusion of tau droplets using dual-trap optical tweezers. Top panel, time-course of the fusion event (brigthfield image) aligned to the laser signal (lower plot) recorded during fusion relaxation (see also movie 1). Combined signal of the two traps are shown in the graph. Data was fit with two exponentials (magenta line). The τ constant (gray rectangle) for 26 fusion events of the fast, initial relaxation was plotted against the characteristic length of the droplet using the geometrical radius (left inlet graph) to extract the ratio of dynamic viscosity to surface tension (slope). Data was fit with a robust linear fit (magenta line). Note the representation of 2 outliners by arrows with their position indicated. **(D)** Internal rearrangement of tau drops. Time-course of Fluorescent Recovery After Photo-bleaching (FRAP) after internal photo-bleaching of tau drops. LUT, FIJI 16 colors. See also movie 2. Bottom panel, plot of the recovery in the photo-bleached area. Values shown are the mean ± SD, n=17. Time 0 indicates the time for the photo-bleaching. Values were normalized to the first time point before photo-bleaching. **(E)** Tau drop deformation by shear flow. Snapshots of movie 3. Shear flow applied from top-left to bottom-right of the image.

**Fig. 2.**
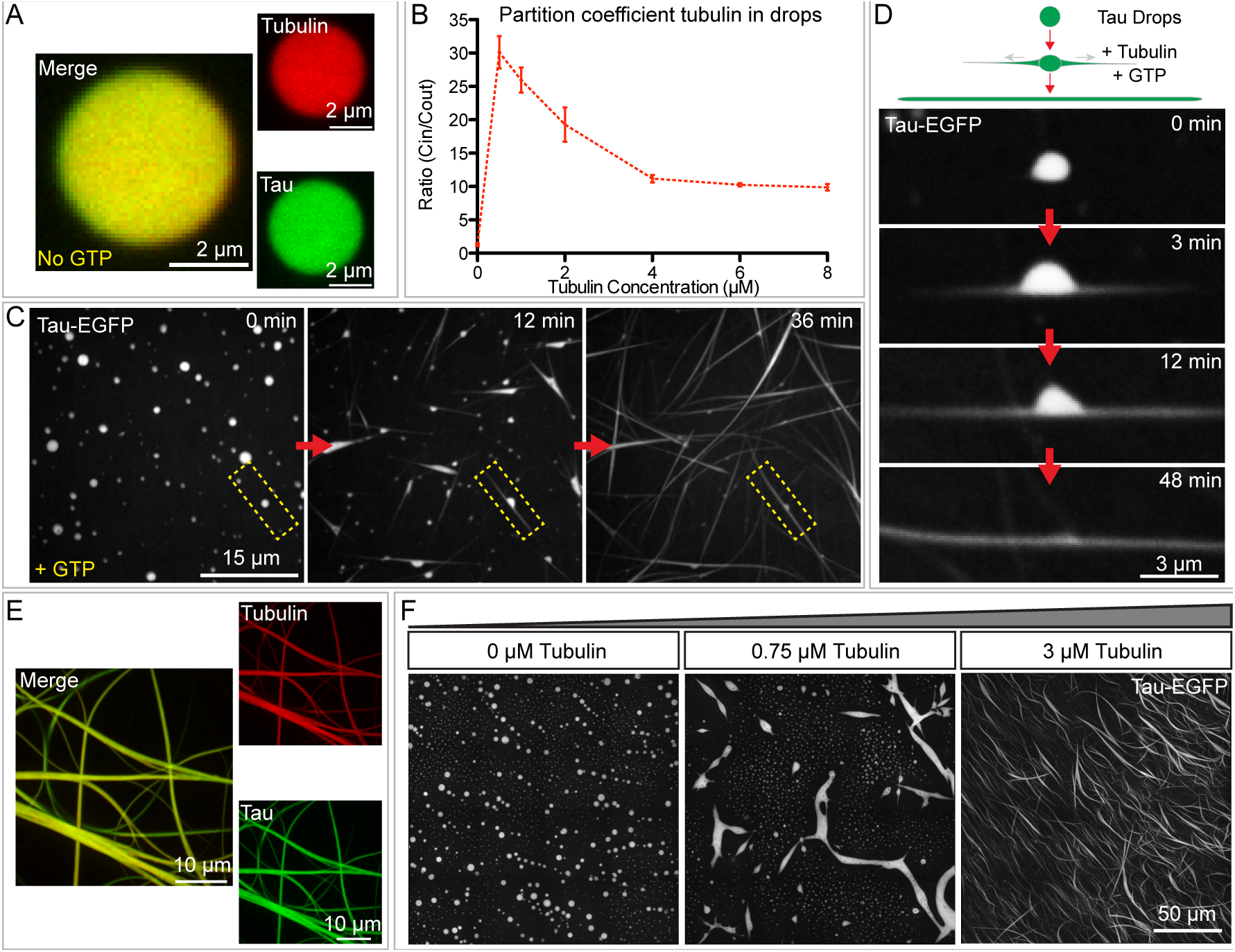
Tau drops concentrate tubulin and polymerize tau-encapsulated microtubule bundles. **(A)** Partition of tubulin into tau drops. Merged and single channel fluorescent images of tau drops (green) with incorporated tubulin dimers (red) 5 minutes after mixing. A final concentration of 5 μΜ rhodamine-labeled tubulin was added to tau drops. Tau drops were formed with 25 μΜ tau-EGFP in 1x BRB80 (pH 6.9) with 1 mM DTT and 10% dextran in all experiments with tubulin if not otherwise mentioned. **(B)** Tubulin partition coefficient quantified by the ratio of the mean intensity inside the drops to the mean intensity in the surrounding bulk media at different concentrations of overall tubulin (no GTP added). Values shown are the mean ± SD, n=16 image stacks, 50 images per stack, mean of all drops in the stack. The partition coefficient of tubulin into tau drops was above 10 in all concentration of tubulin tested, see also figure S2. **(C)** Tau drops deformation by internal microtubule bundles polymerization. 5 μΜ rhodamine-tubulin was added to tau drops together with 1 mM GTP. Immediately after addition of tubulin the drops deformed. See also movies 4. All experiments in this paper were performed at room temperature. **(D)** Detail from previous panel of a single drop deformation upon addition of tubulin and GTP. Drops deformed bidirectionally due to the polymerization of internal nucleated microtubule bundles. Drops redistributed their volume along the growing microtubule bundles. See also movies 5. **(E)** Tau and tubulin co-localization in tau-encapsulated bundles formed from the deforming drops. Single channel and merged fluorescent maximum projection images are shown. The concentrations and conditions used are the same as in previous panels. **(F)** Nucleation of tau-encapsulated bundles at low tubulin levels. 0.75 μΜ overall tubulin concentration is sufficient to observe the deformation of some drops by the internally nucleated bundles, 15 min after addition at room temperature. See also supplementary figure S4. Stitching of 16 maximum projection images are shown in each panel. Rhodamine-labeled tubulin was added at the indicated concentration to tau drops.

**Fig. 3.**
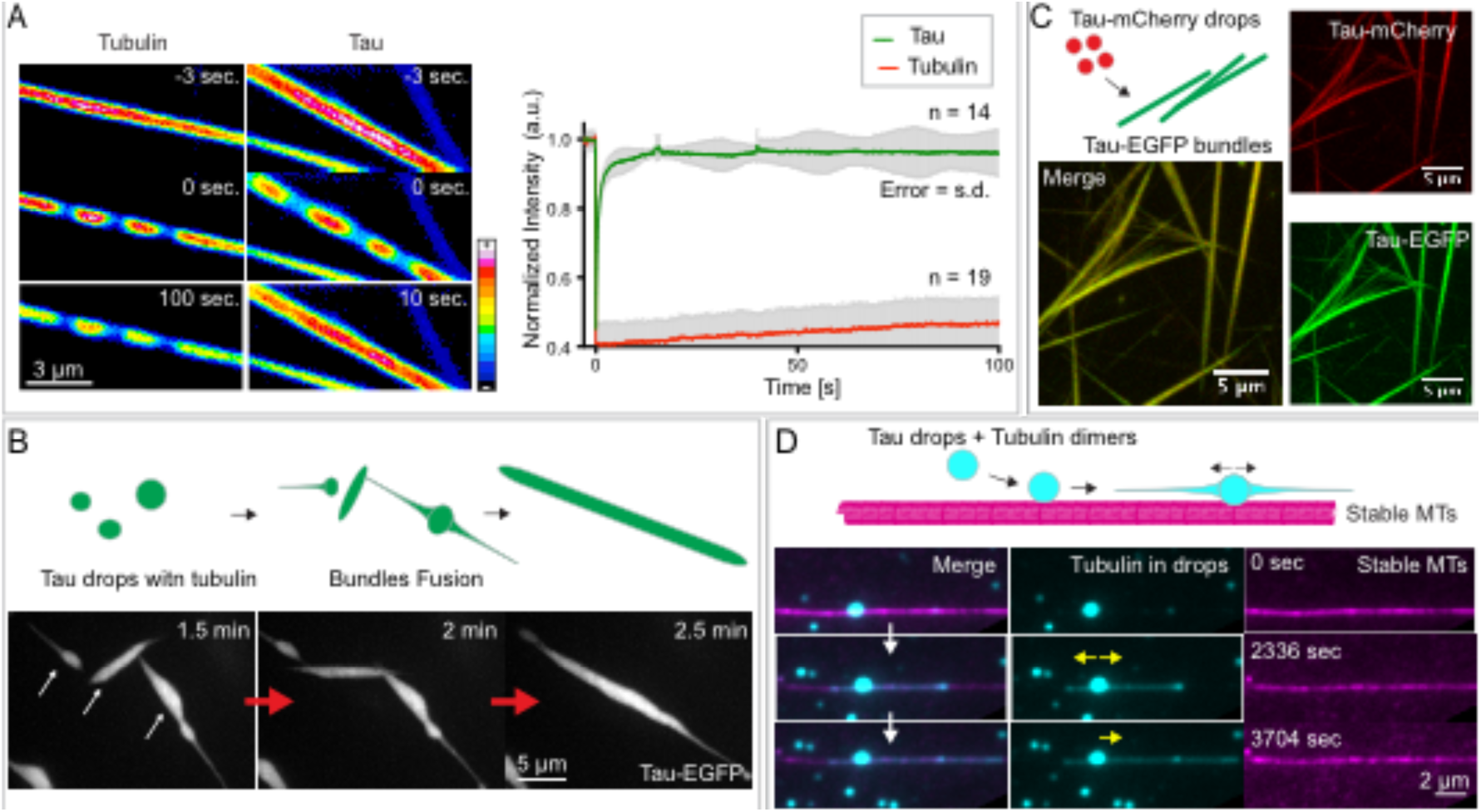
Tau in tau-encapsulated microtubule bundles has liquid-like properties. **(A)** Internal rearrangement of tubulin and tau in tau-encapsulated microtubules bundles. Left panel, time-course fluorescent images of FRAP for both tubulin (left row) and tau (right row) in bundles formed from tau drops. The recovery of 3 photo-bleached regions in the bundles is shown. LUT, FIJI 16 colors. See also movie 7. Right panel, quantification of the recovery for both tau (green) and tubulin (red) in bundles. Values shown are the mean ± SD, n=14 and 19 respectively. Bundles were formed from tau drops with protein concentrations and buffer conditions as mentioned in previous figure. **(B)** Tau encapsulated microtubule bundles fuse over-time. Time-course for the fusion of three bundles (see also movie 8). Maximum projection fluorescent images of tau-EGFP are shown. Bundles were formed as mentioned before. **(C)** Fusion of tau-mCherry drops added to preformed tau-EGFP bundles. Bundles were formed from tau-EGFP drops as before but adding 5 μΜ of unlabelled tubulin together with 1 mM GTP. Tau-mCherry drops were formed with 25 μΜ of tau-mCherry in same buffer conditions and added 1:2 to the bundles. **(D)** Guided growth of microtubules polymerizing from drops along preexisting microtubules. See also movie 10. Tau drops were formed as in previous panels, 5 μΜ Cy5-tubulin together with 1 mM of GTP was added to the drops and flushed into a flow chamber containing immobilized and stable rhodamine-labeled microtubules. Microtubules growing from drops (cyan, Cy5-tubulin) grow in the direction of the attached preassembled microtubule (magenta). Microtubules were stabilized with taxol and GMP-CPP.

**Fig. 4.**
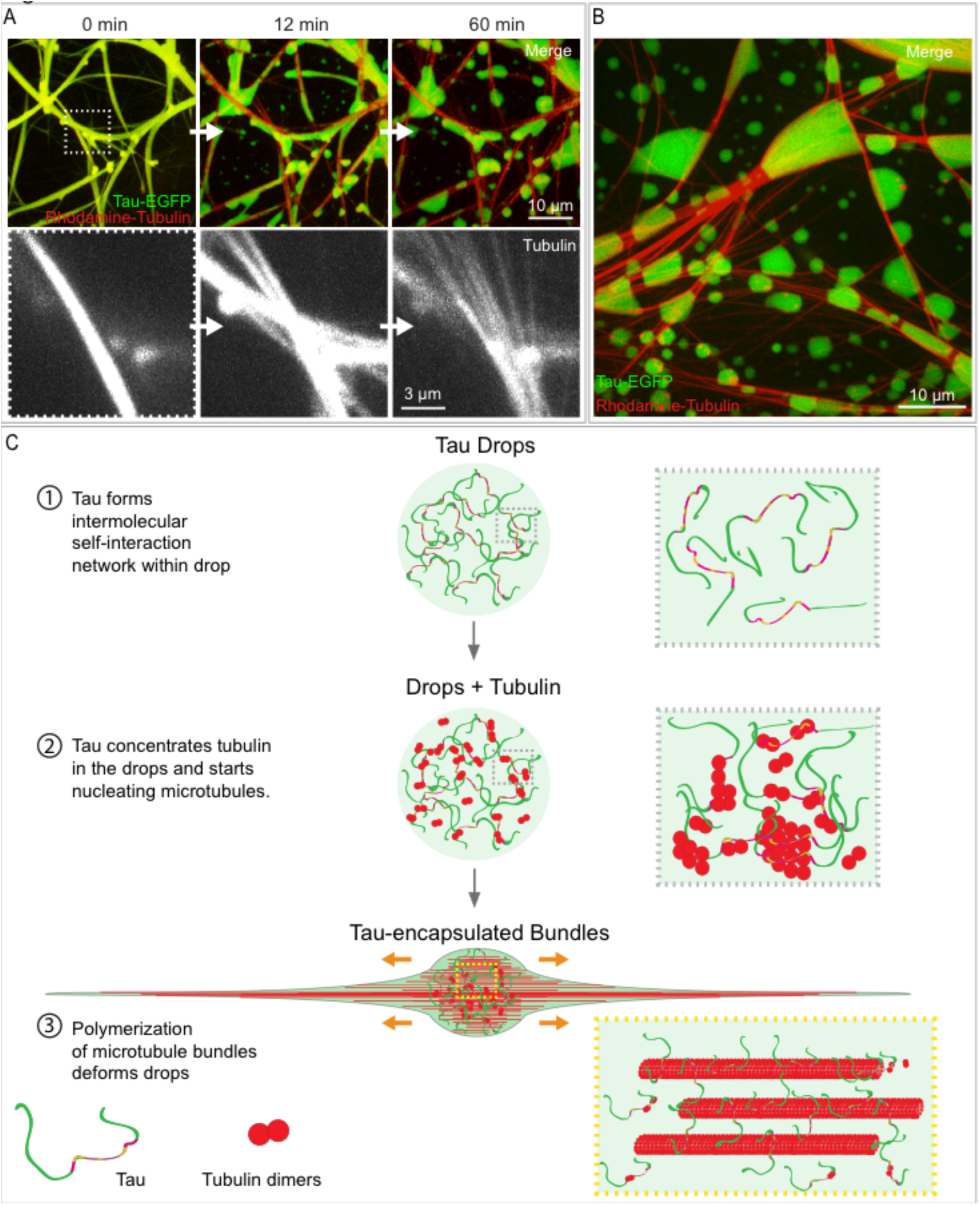
Tau encapsulation maintains microtubule bundles. **(A)** Heparin addition to tau-encapsulated bundles. Upper panels, time-course of tau dewetting from microtubules bundles, its reshaping back into drops and the simultaneous debundling of microtubules after addition of heparin. Tau-EGFP shown in green and rhodamine-tubulin in red. See also movie 11. Lower panel, detail of the microtubules debundling. 200 μg/ml of heparin was added to bundles formed with 5 μΜ rhodamine-tubulin and 1 mM GTP added to tau drops (protein concentration and buffer conditions as in previous figures). Maximum projection images shown. **(B)** Approximately one hour after heparin addition, tau is completely reshaped into both free and microtubule-attached drops. Conditions used are as in previous panel. **(C)** Model of tau-encapsulated microtubule bundles assembly. (1) Tau’s intrinsically disordered properties enable its phase separation in a crowded environment. (2) Tubulin dimers get concentrated inside tau drops allowing microtubule nucleation. (3) Microtubule bundles growing within drops, deform it into a rod-like shape, remaining encapsulated by liquid tau. Tau simultaneous binding to multiple tubulin dimers by its four tubulin binding repeats, together with its intrinsically disordered arms may be important for tubulin concentration and microtubule bundles nucleation and stabilization.

## EXPERIMENTAL PROCEDURES

### Prediction of tau protein disorder distribution and low complexity domains

To predict the disorder degree along human tau protein (splice variant htau441) PONDR program was employed (www.disprot.org/index.php). PONDR-FIT (P-FIT) and VL3 algorithms are shown. These algorithms are generally used to predict the disorder tendency of proteins that are experimentally known to be disordered (Obradovic et al., 2003; Xue et al., 2010). The low complexity domains in htau411 were predicted with SEG program (http://mendel.imp.ac.at/METHODS/seg.server.html, Wootton, 1994). LCD1 region comprises amino acids 129 to 153 and LCD2 amino acids 172 to 223. The level of stringency used was SEG 12 2.2 2.5.

### Constructs

The cDNA encoding for htau441 (also named 2N4R) was a gift from Peter Klein [tau/pET29b, Addgene plasmid number # 16316, (Hedgepeth et al., 1997)]. Htau441 cDNA was amplified by PCR from this vector using the primers shown in Table S1 and cloned into a baculoviral expression vector (pOCC series). Details of constructs used are listed in Table S1.

His tag cleaved version of htau441-EGFP-PS-6xhis and htau441-mCherry-PS-6xhis were used in this study. Untagged version of Tau (C-terminal 6xhis cleaved) and N-terminal EGFP tagged version of Tau were also purified as controls (htau441-PS-6xhis and 6xhis-PS-EGFP-htau441) to assess is capability to form drops and bundles. In both cases drops and bundles formed. A linker was always present between Tau and the fluorescent tag. All constructs generated were confirmed by digestion and sequencing. PS refers to PreScission protease recognizing sequence.

### Recombinant protein expression and purification

#### Tau variants

All tau variants were purified from insect cells using the baculovirus infection system. Recombinant baculovirus for each construct were produced according to Woodruff and Hyman (2015). Sf9 insect cells (log phase, 1 million cells/ml, Expression system, Cat#94-001F) were infected with 5 ml of P2 baculovirus stock, incubated at 27°C with moderate shaking and harvested 72 hours post infection. Cells were harvest by centrifugation at 700g for 8 min and resuspended in resuspension buffer [25 mM Hepes, 150mM KCl, 20 mM imidazole (pH 7.4) with 1 mM DTT, 1 mM PMSF (Sigma) and 1x Protease inhibitors cocktail (Calbiochem, Type III)]. Cells were lysate using an Emulsiflex (Emulsiflex-C5, Avestin). Cell lysate was spin at 35000 rpm for 45 minutes and the supernatant was collected. The lysate was filtered through a 0.45 μm filter and incubated with Ni-NTA agarose resin (QIAGEN) HiTrap for 1 hour. Disposable gravity columns (20 mL, Biorad) were used to collect and wash the beads. Columns were washed with 10 columns volume with wash buffer (25 mM Hepes, 150 mM KCl, 30 mM imidazole, 1 mM DTT, pH 7.4) and eluted with elution buffer (same buffer with 250 mM imidazole). 6xhis tag was removed by treatment with PreScission protease (3C HRV protease, 1:100, 1μg enzyme/100 μg of protein, overnight at 4°C). Imidazole was removed by dialysis (slide-a-lyzer with 20KDa cut off, overnight at 4°C). His tag cleave was performed simultaneously with the dialysis. The protein was further purified by size-exclusion chromatography using a HiLoad 16/60 Superdex 200 column with and ÄKTA Pure chromatography system (GE Healthcare) in 25 mM Hepes, 150mM KCl, 1 mM DTT (pH 7.4). Collected peak fractions were concentrated to 100 μM or 200 μM using Amicon Ultra 30K (Millipore). Protein concentration was measured with a NanoDrop ND-1000 spectrophotometer (Thermo Scientific) at 280 nm absorbance. Proteins were flash-frozen in liquid nitrogen and stored at −80**°**C. All steps in the purification were performed at 4°C. Purified proteins are shown in Fig. S1A.

#### Tubulin

Tubulin was purified from porcine brains as previously described (Gell et al., 2011). Purified tubulin used is shown in figure S1A. Cycled tubulin was dimly labeled with rhodamine or Cy5 fluorescent dyes as previously described (Hyman et al., 1991).

### *In Vitro* assays

#### Drop formation

36 to 50 μM tau-EGFP in 25 mM Hepes, 150 mM KCl (pH 7.4) with freshly added DTT (1 mM) was mixed with 20% dextran (T500, Pharmacosmos) 1:1 to a final concentration of 18 to 25 μM tau-EGFP and 10% dextran. 18 μM of tau-EGFP was the lowest concentration at which we observed drops formation at 10% dextran without any other nucleation factor. 25 μM of tau-EGFP was the final concentration of tau used along this study if not otherwise mentioned. T500 dextran was also used throughout this study. Tau drops were also observed using 10% of other crowding agents like polyethylene glycol (PEG-8000, Sigma) or Ficoll-400 (Sigma). In all the experiments involving tubulin, 1x BRB80 (80 mM PIPES, 1 mM EGTA, 1 mM MgCl_2_, pH 6.9) was used to dilute the protein and the dextran if not otherwise mentioned.

#### Droplets fusion with optical tweezer

Controlled drop fusion experiments were conducted in a custom-built dual trap optical tweezer microscope with two movable traps (Jahnel et al., 2011; see also Patel et al., 2015). Tau droplets were formed with 25 μM tau-EGFP in 25 mM Hepes, 150 mM KCl (pH 7.4) with 10% dextran and 1 mM fresh DTT. 5 μl of this mix were placed in a static flow chamber (coverslip – double-sided tape – coverslip sandwich). Droplets could be trapped at minimal laser intensity (laser power at focal position < 50 mW to prevent heating artifacts) due to a mismatch in the index of refraction between the droplets and the surrounding media. Keeping one optical trap (T1) stationary, the other optical trap was moved until droplets were brought into contact, after which droplet fusion was recorded with 1 ms temporal resolution (1 kHz). The combined laser signal showed two characteristic phases: a fast rise potentially driven by surface tension, and a slower relaxation, governed by viscosity. To quantify the dynamics of tau droplet fusion we describe the process phenomenologically as a two state exponential relaxation process with the following formula for the laser signal f(t) (magenta fit to time series in Fig. 1c):

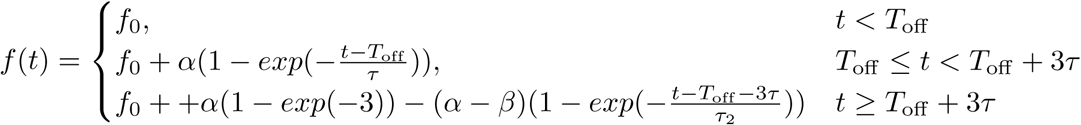
where f_0_ is the signal base line, T_off_ is the starting time of the fusion process, τ and τ_2_ are the fast and slow relaxation times, and α and β are the plateau values of the fast and slow relaxation regimes, respectively.

Note that the fusion signal could not be well described as a single exponential relaxation process, which was sufficient for other protein liquids studied with the same method (Patel et al., 2015; Saha et al., 2016). This suggests the onset of inertial effects, a transition often challenging to study in more conventional liquids (Eggers et al., 1999). Plotting the time constant, τ, for the fast initial relaxation process against the characteristic length scale of the droplets (geometric radius, r = √ (r1 * r2)) we estimate the ratio of dynamic viscosity to surface tension to be 13 ± 2 ms / μm (magenta fit ± 95% confidence interval in left inlet of Fig 1c).

#### Wetting

A shear flow was generated by pressing one side of a flow channel generated with silicone grease and two parallel double size tape in between a coverslips and a glass slide. Images were acquired using an upright spinning microscope (Leica, DM6000 B) with spinning disc head (Perkin Elmer) using a 63x glicerol-immersion objective (UPlanSApochromat, NA 1.3).

#### Fluorescence Recovery After Photo-bleaching (FRAP)

FRAP was performed as previously described (Brangwynne et al., 2009) in a spinning-disc confocal imaging system (inverted Olympus microscope with a Yokogawa CSU-X1 scan head) fitted with diode-pumped solid-state lasers [488 nm Coherent Sapphire (50mW) and 561 nm Cobolt Jive (50mW)] using a 100x objective (UPlanSApochromat 100x NA 1.4 oil-immersion) and Andor iXon DU-897 BV back illuminated EMCCD camera. Single confocal plane was imaged over time for all FRAP experiments. A roi of ∼1,07 μm^2^ was photo-bleached. Image were acquired every 100 milliseconds. 30 and 1000 images were recorder before and after photo-bleaching, respectively.

#### Frap analysis

FRAP Calculator macro generated by Robert Bagnell was used to analyze the bleached regions (https://www.med.unc.edu/microscopy/resources/imagej-plugins-and-macros/frap-calculator-macro). The plugin corrects data for photo-bleaching due to the imaging process. Values were normalized to the intensity prior to photo-bleaching.

#### Microtubule bundles formation

Tau drops were formed with 25 μΜ tau-EGFP and 10% dextran in 1xBRB80 with 1 mM DTT. Cycled Tubulin (unlabeled or rhodamine-labeled) was added to the drops at the concentration mentioned for each experiment together with 1 mM GTP. Drops deformation and bundle formation equally occurred in 25 mM Hepes, 150 mM KCl, pH 7.4 when tubulin and GTP was added. All experiments were carried out at room temperature.

#### Microscopy

Drops and bundles were imaged within flow chambers generated by attaching double-sided tape (∼150 μm thick) to a glass slide and sealed with a microscope slide after addition of 5 or 10 μl of sample. For conditions where reagents were added during imaging or just prior to imaging, MatTek-glass bottom dish were used. To avoid evaporation in these cases, mineral oil (Cat# 330779, Sigma) was added on top of the sample.

In all experiments oxigen scavanger mix was added to assay buffer at the following concentrations; 40 mM D-Glucose (Cat# G7528, Sigma), 56 μg/ml, Glucose oxidase (Aspergillus niger, Cat# 22778, SERVA electrophoresis GmBH), and 11 μg/ml Catalase (Cat# C40, Sigma).

#### Interaction with single microtubules and TIRF imaging

Saturation experiments on single microtubules (Fig. S5A,B), zippering of tau-coated microtubules (Fig. S5C and movie 9) and drops interaction with single stable microtubules (FIG. 3D) were imaged with TIRF microcopy. Microtubules and flow chambers for these experiments were prepared as described previously (Fink et al., 2009). Microtubules and tau-EGFP molecules were visualized using an inverted fluorescence microscope (Axio Observer Z3, Carl Zeiss) with a Zeiss 63x oil immersion 1.46 NA TIRF objective and a build-in 1.6x optovar in combination with an Andor iXon 897 (Andor Technology) EMCCD camera controlled by Metamorph (Molecular Devices Corporation). A Lumen 200 metal arc lamp (Prior Scientific Instruments) for excitation in epi-fluorescence mode and a 488 nm, 50 mW, diode solid-state laser (Vortran) for TIRF-illumination in combination with GFP and TRITC filters (Chroma Technology) were used.

#### Saturation experiment

Tau-EGFP was added to surface-immobilized, rhodamine-labeled GMP-CPP stabilized microtubules in increasing concentrations ranging from 1 nM to 20 μM in presence or absence of 10 % dextran in assay buffer (20 mM Hepes pH 7.4, 75 mM KCl, 1 mM EGTA, 1 mM MgCl2, 10 mM DTT, 0.5 mg/ml casein, 20 mM D-glucose, 110 μg/ml glucose oxidase, and 20 μg/ml catalase). To estimate the number of tau molecules per μm of microtubule length, tau-EGFP integrated intensities along a microtubule were divided by the average intensity of single molecules of tau-EGFP in images acquired under identical imaging conditions, and then divided by the length of the microtubule.

#### Zippering of single microtubules by tau

25 μM tau-EGFP in assay buffer was added to surface-immobilized GMP-CPP stabilized microtubules. When two microtubules where laying close to each other, or crossed at very shallow angles, the microtubules zippered up.

#### Interaction of tau drops with preexisting microtubules (guidance)

Double stabilized rhodamine-labelled microtubules templates (GMPCPP and 10 mM paclitaxel, 1x BRB80, pH 6.9) were injected into flow chamber and bound to surface-immobilized tubulin anti-bodies. After rinsing the chamber with assay buffer (25 mM HEPES 10 mM DTT, 0.5 mg/ml casein, 10 mM paclitaxel, 0.1% Tween, 40 mM D-glucose, 110 μg/ml glucose oxidase, and 20 μg/ml catalase), tau drops with overall concentration of 5 μM Cy5-labeled tubulin dimers was flushed into the channel.

In all TIRF image experiments oxigen scavanger mix was used.

#### Heparin treatment

Heparin sodium salt from porcine intestinal mucosa (Cat# H3393, Sigma) was added to tau-encapsulated microtubules bundles at a final concentration of 200 μg/ml.

#### Image analysis

Images were analyzed using Image J (FIJI). For tubulin concentration in tau drops and tau drops nucleation by tubulin a stack image (50 stacks per image) were segmented using a signal threshold. Camera background was subtracted before calculating mean fluorescence intensities and partition coefficient (ratio mean intensity in the drop versus mean intensity in the bulk solution). The script can be found in the supplementary file S1.

#### Concentration of tubulin in drops

Mean fluorescent intensity of rhodamine-tubulin was measured from an image z-stack at a final total concentration of 1 μM tubulin for each condition. The mean intensity in any of the conditions without drops (see fig, S2B) was comparable to the one in the bulk medium in conditions were drops were present. To estimate the concentration of tubulin in the drops we assumed that the relation between intensity and concentration was linear and compared the intensity in the drop to the mean intensity in the conditions with tubulin alone. At 20 minutes after formation the concentration was estimated to be ∼ 25 μM.

**Table S1.**
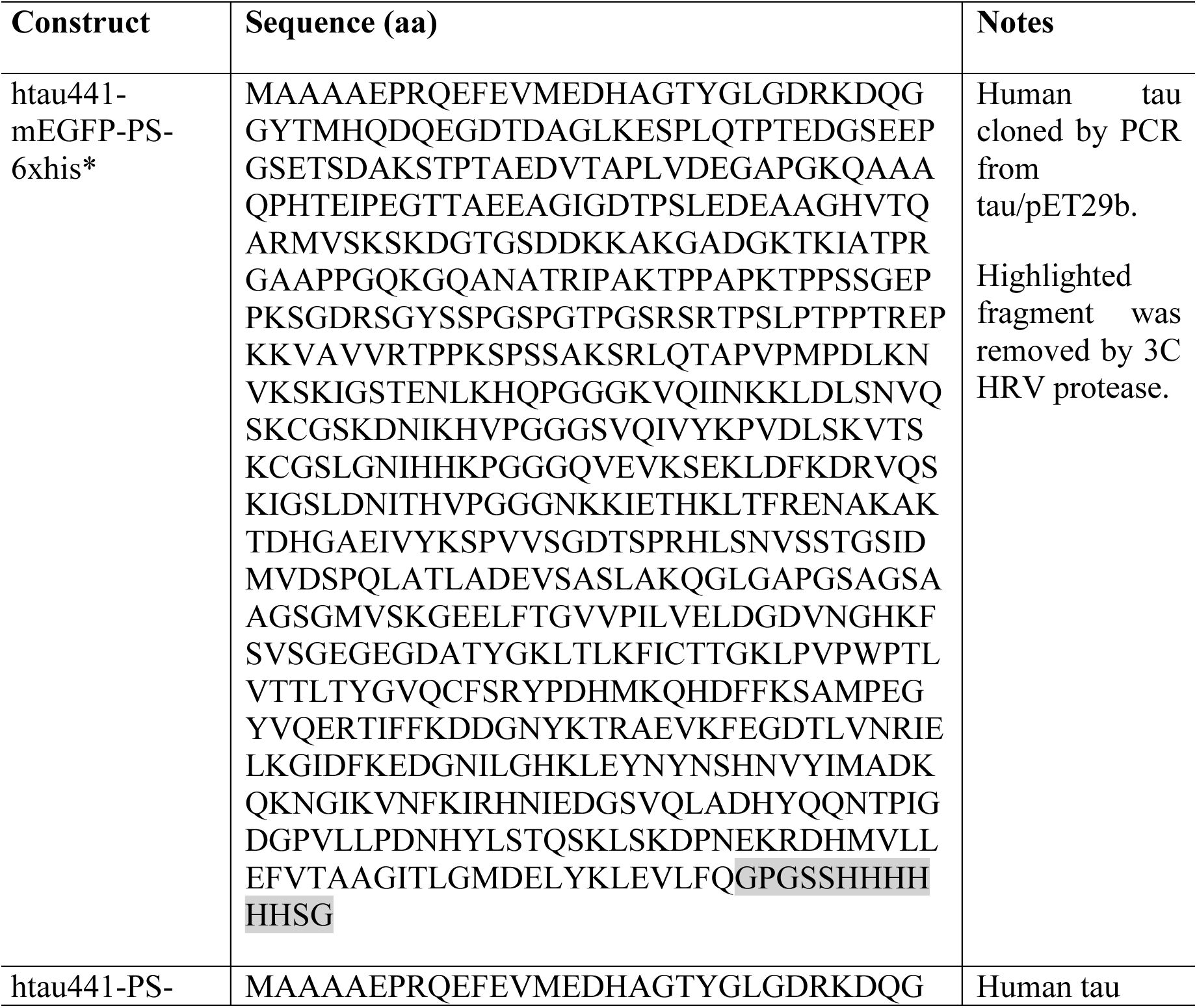
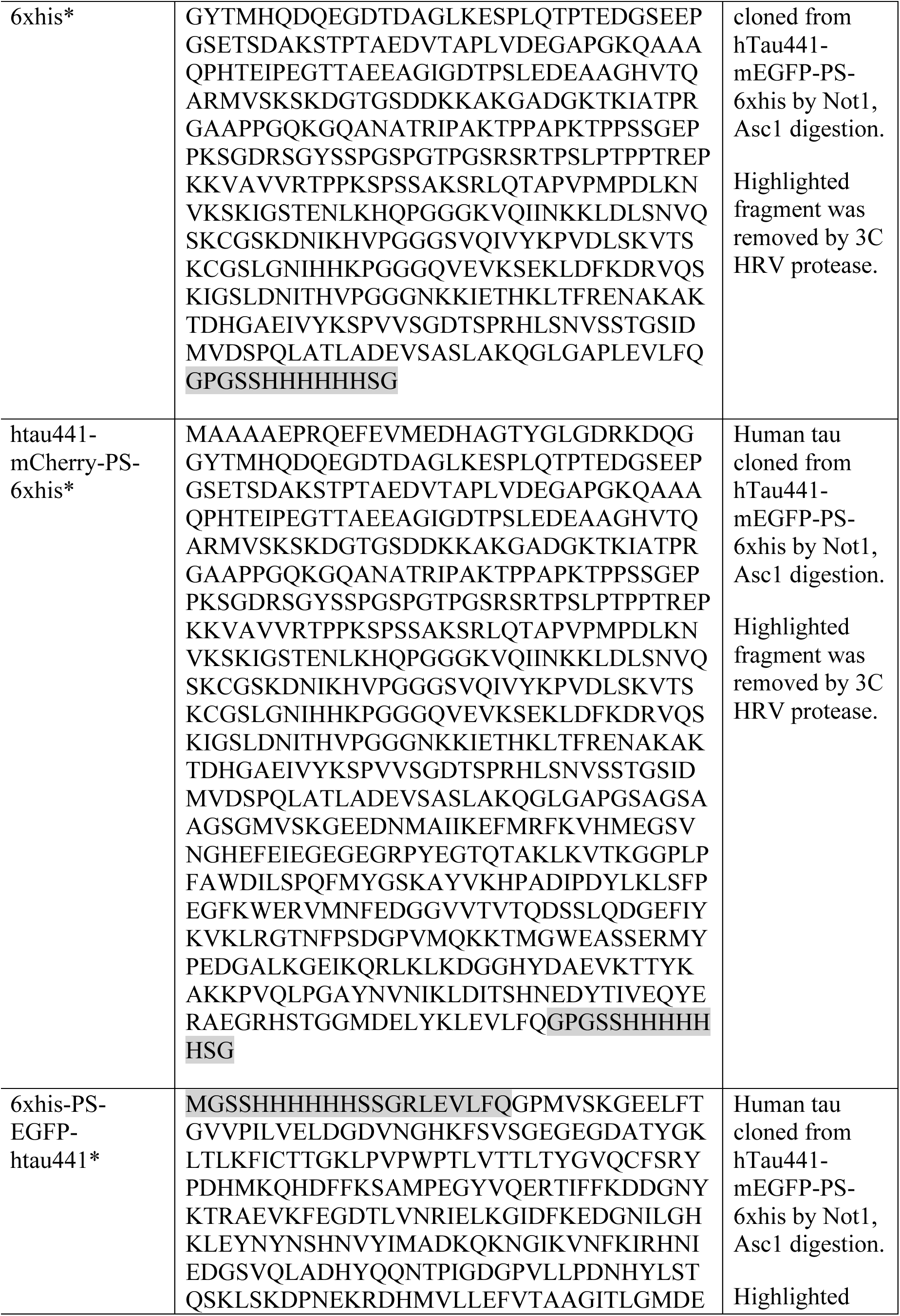
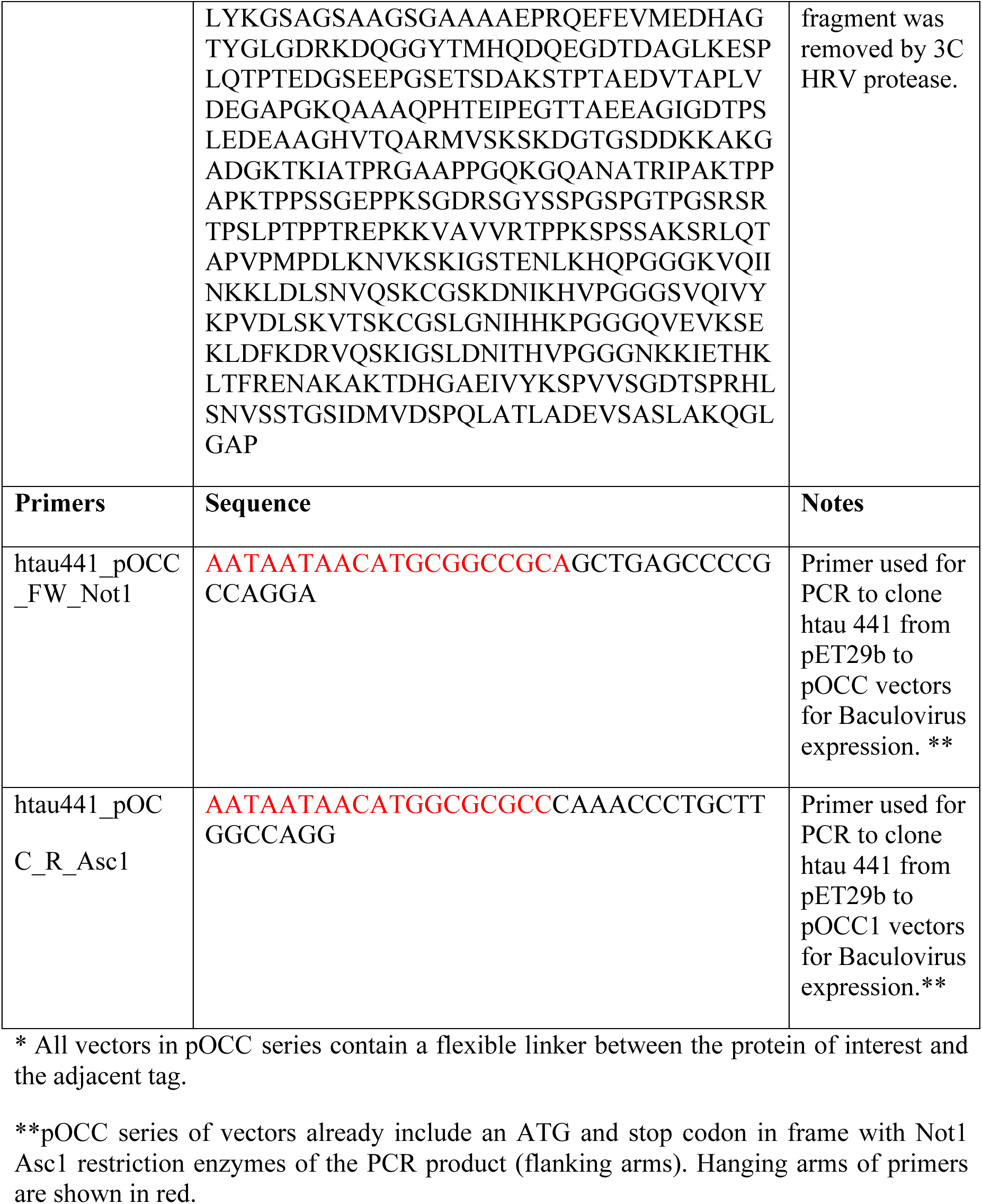

## Acknowledgements

We thank David Drechsel, Jan Brugués, Gaia Pigino, Simon Alberti, Titus Franzmann, Avinash Patel and Mark Leaver for comments and discussions of earlier versions of this manuscript. We thank the Protein Expression and Purification facility at MPI-CBG (Dresden) especially David Drechsel, Barbara Borgonovo, Regis Lemaitre and Martine Ruer for help with protein purification. The Light Microscopy facility at MPI-CBG (Dresden) for help with image acquisition. Benoit Lombardot from the Bio-Image informatics, Scientific Computing Facility at the MPI-CBG (Dresden) for developing the script to segment and analyze drops number and partition coefficients. Andrei Pozniakovsky and Aliona Bogdanova for help with cloning DNA constructs. Avinash Patel, Titus Franzmann, Shamba Saha, Jie Wang, Mark Leaver and other members of Hyman, Diez and Alberti laboratories for suggestions and help in *in vitro* experiments.

## SUPPLEMENTAL INFORMATION

**Fig. S1.**
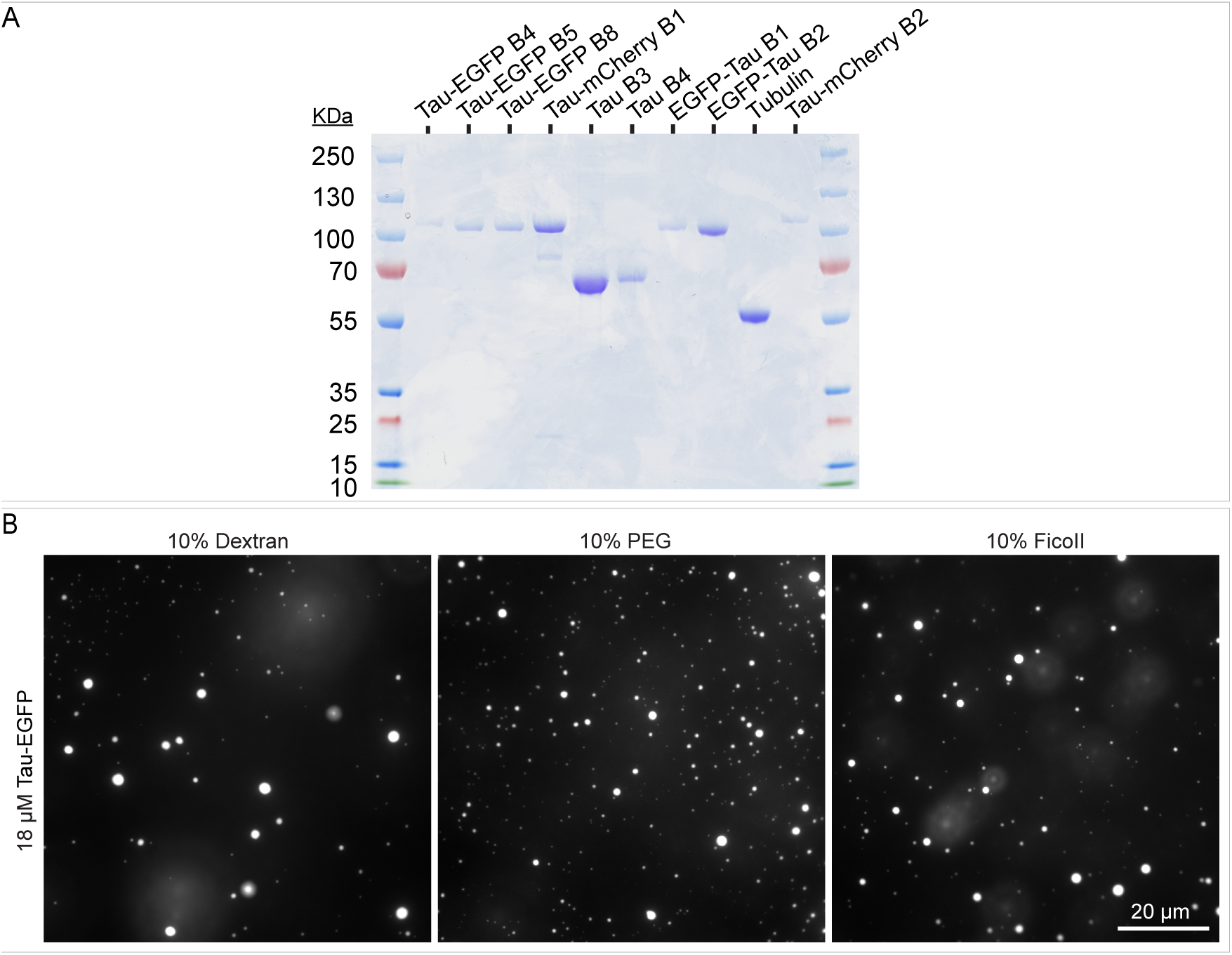
*In vitro* tau drops formation in the presence of different crowding agents. **(A)** SDS-PAGE of the purified proteins used in this study (stained with Coomassie blue). B refers to protein batch. **(B)** *In vitro* formation of tau drops in the presence of different crowding agents. Drops were formed with 18 μΜ of tau-EGFP, 25 mM Hepes, 150 mM KCl, 1 mM DTT, pH 7.4 in the presence of 10% of the indicated crowding agents.

**Fig. S2.**
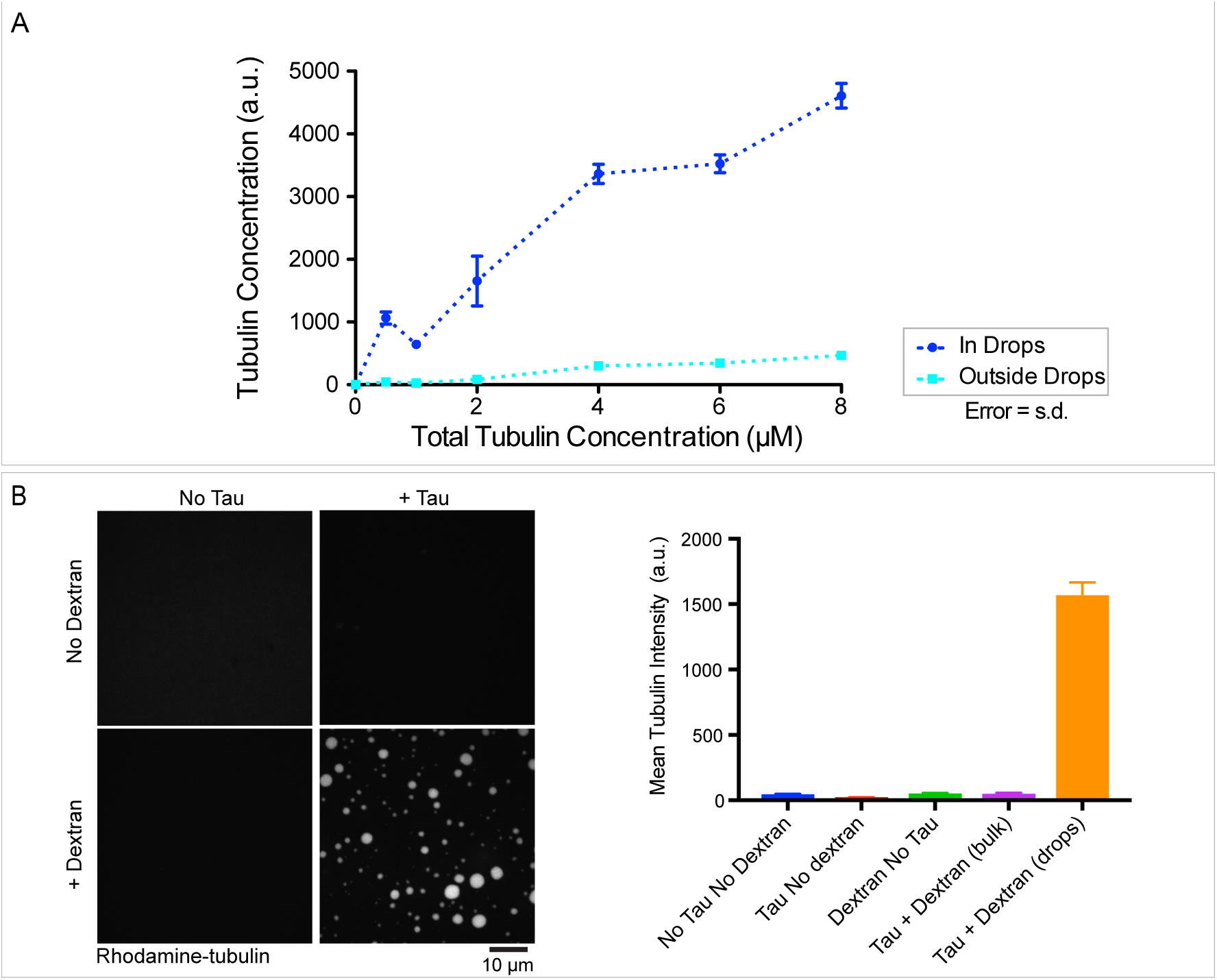
Tubulin partition into tau drops. **(A)** Tubulin concentration inside tau drops (dark blue) and in the surrounding bulk media (pale blue) quantified by mean intensity at different concentrations of total levels of tubulin (in absence of GTP). Values shown are the mean ± SD. n=16 image stacks, 50 images per stack. **(B)** Estimation of tubulin concentration in drops. rhodamine-labeled tubulin mean intensity at 1 μΜ of total level of tubulin in conditions without drops (tubulin or tubulin plus dextran or tau) or in the presence of drops (tau plus dextran and tubulin). Proteins were dissolved in 1x BRB80, with 1 mM DTT and 10% dextran or 25 μΜ of tau when indicated. Assuming a linear relation between intensity and concentration for fluorescently-labeled tubulin, the estimated tubulin concentration in the drops is higher than 20 μΜ 20 min after the addition of 1 μM overall concentration of tubulin at room temperature (see M&M for details in the estimation). All experiments were carried out at room temperature.

**Fig. S3.**
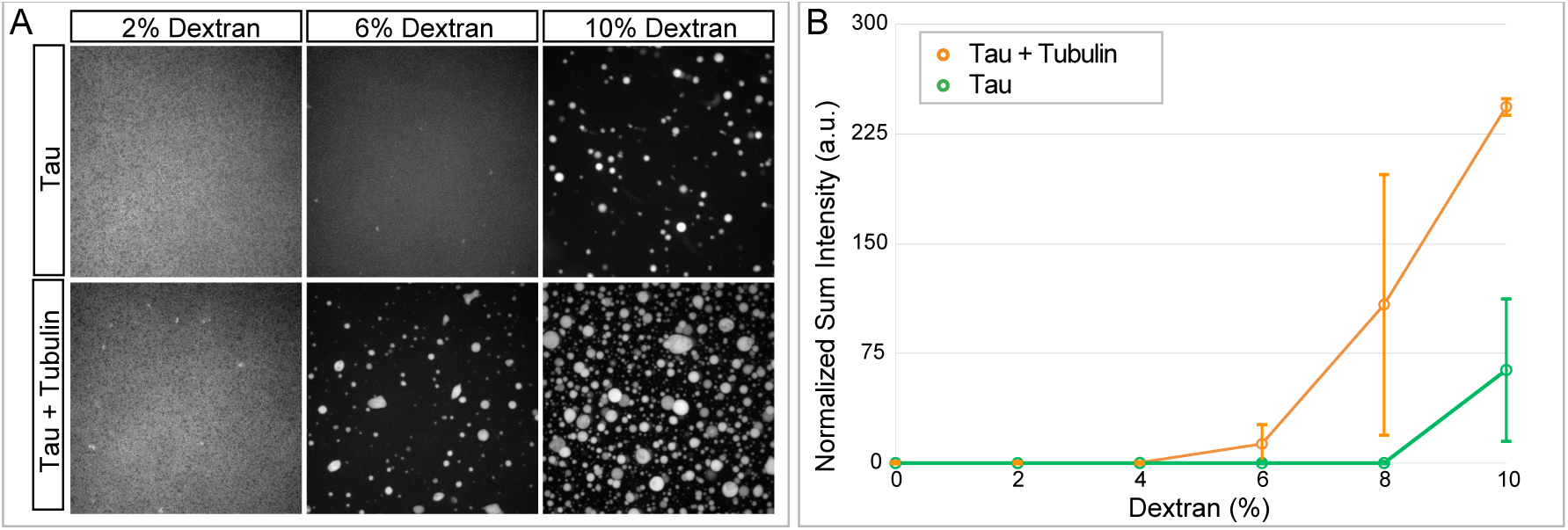
Tubulin enhances tau drops nucleation. **(A)** Representative images of tau drops formation at different concentrations of dextran in the presence or absence of 8 μΜ unlabelled tubulin (in absence of GTP). **(B)** Normalized sum intensity (sum of intensity in all drops detected in a z-stack) at different concentration of dextran and in the presence or absence of 8 μΜ unlabelled tubulin (in absence of GTP). Values shown are the mean ± SD, n=3, 16 images stack of 50 images per stack were analyzed for each replica. Drops were formed with 25 μΜ of tau-EGFP in 1x BRB80 with the amount of dextran indicated and 1 mM DTT.

**Fig. S4.**
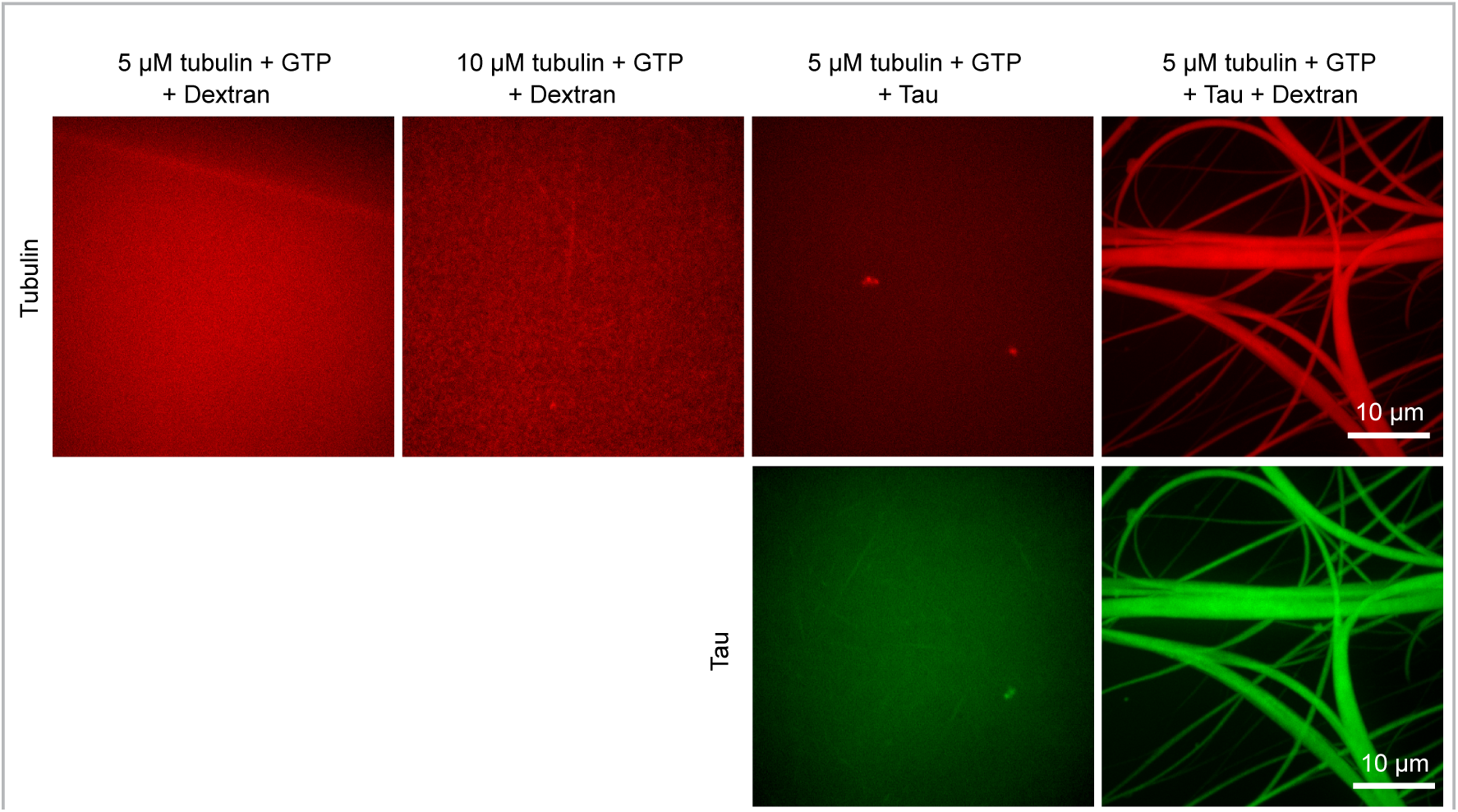
Tau drops are needed for tubulin concentration and bundles polymerization. Upper panel, rhodamine-tubulin. Lower panel, tau-EGFP. Neither, 5 or 10 μM of tubulin in the presence of 10% dextran (room temperature) is sufficient to polymerize microtubules (first and second row). 5 μΜ of tubulin in the presence of tau (no dextran, no drops, room temperature) allows the polymerization of single small microtubules (third row). See also movie 6. In the presence of drops (tau plus dextran), 5 μΜ tubulin is sufficient for the polymerization of long tau-encapsulated bundles of microtubules. 25 μΜ of tau-EGFP in 1x BRB80 with the indicated amount of rhodamine-tubulin and 1 mM GTP plus 1 mM DTT were used. As mentioned, all experiments were carried out at room temperature.

**Fig. S5.**
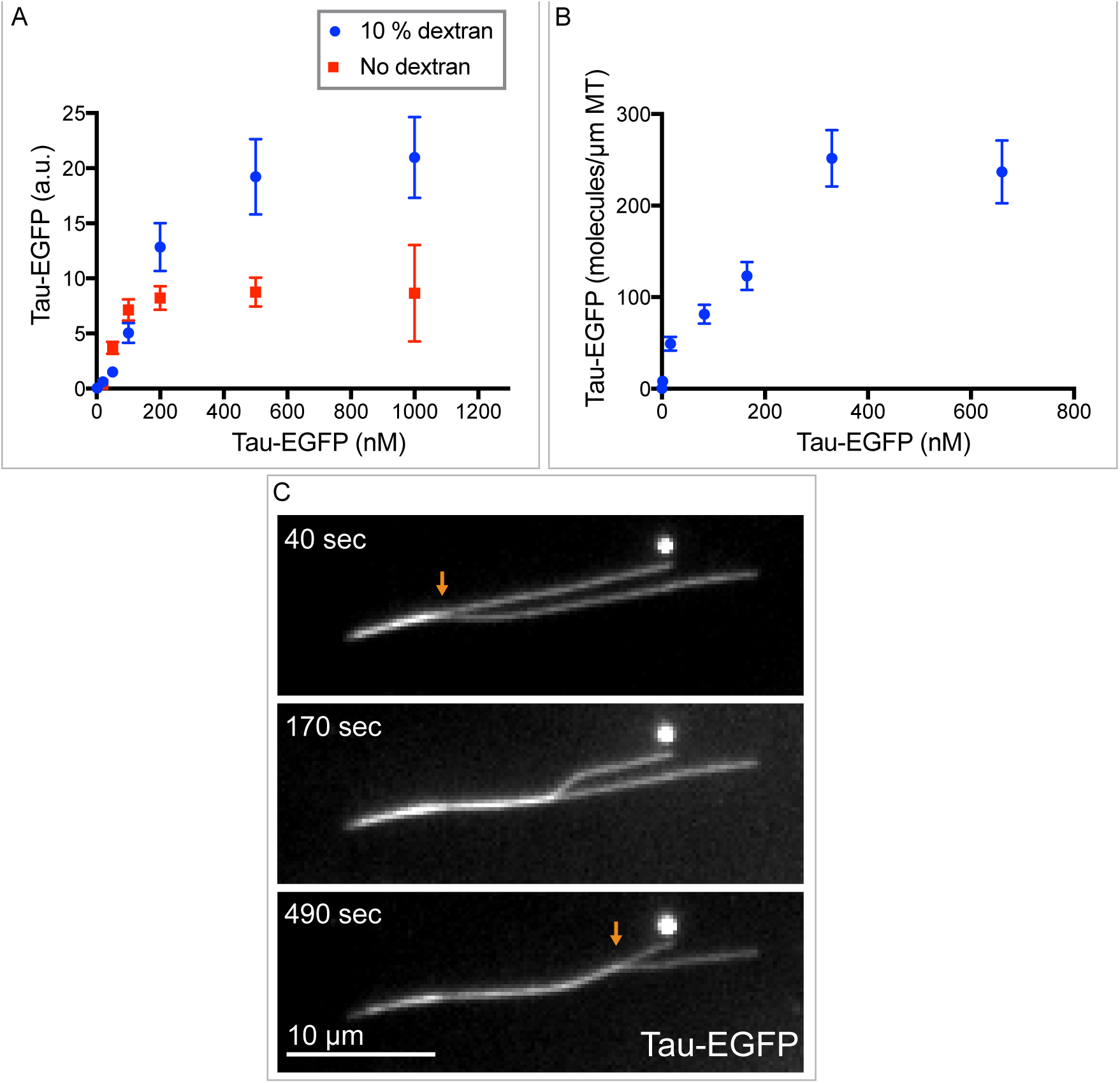
Tau interaction with preassembled stabilized microtubules. **(A)** Amount of tau binding to single stabilized microtubules (GMP-CPP stabilized). Tau shows saturable binding to single preassembled stabilized microtubules both in the presence or absence of dextran (drops or no drops, respectively). Plot shows the signal of tau on immobilized microtubules at different concentrations of soluble tau. Data shown are Mean ± SD, n = 12. **(B)** Estimation of the number of tau molecules binding per μm of stabilized microtubule. Tau single molecule intensity was used to normalize the number of molecules per μm of microtubule. Mean ± SD are shown, n = 7. The experimental conditions are as in (A). **(C)** Zippering of two stable microtubules coated with tau. See also movie 9. A flow cell containing immobilized, stable microtubules, was flushed with a solution of 25 μΜ of tau-EGFP in 10% dextran in BRB80/Hepes buffer.

**Fig. S6.**
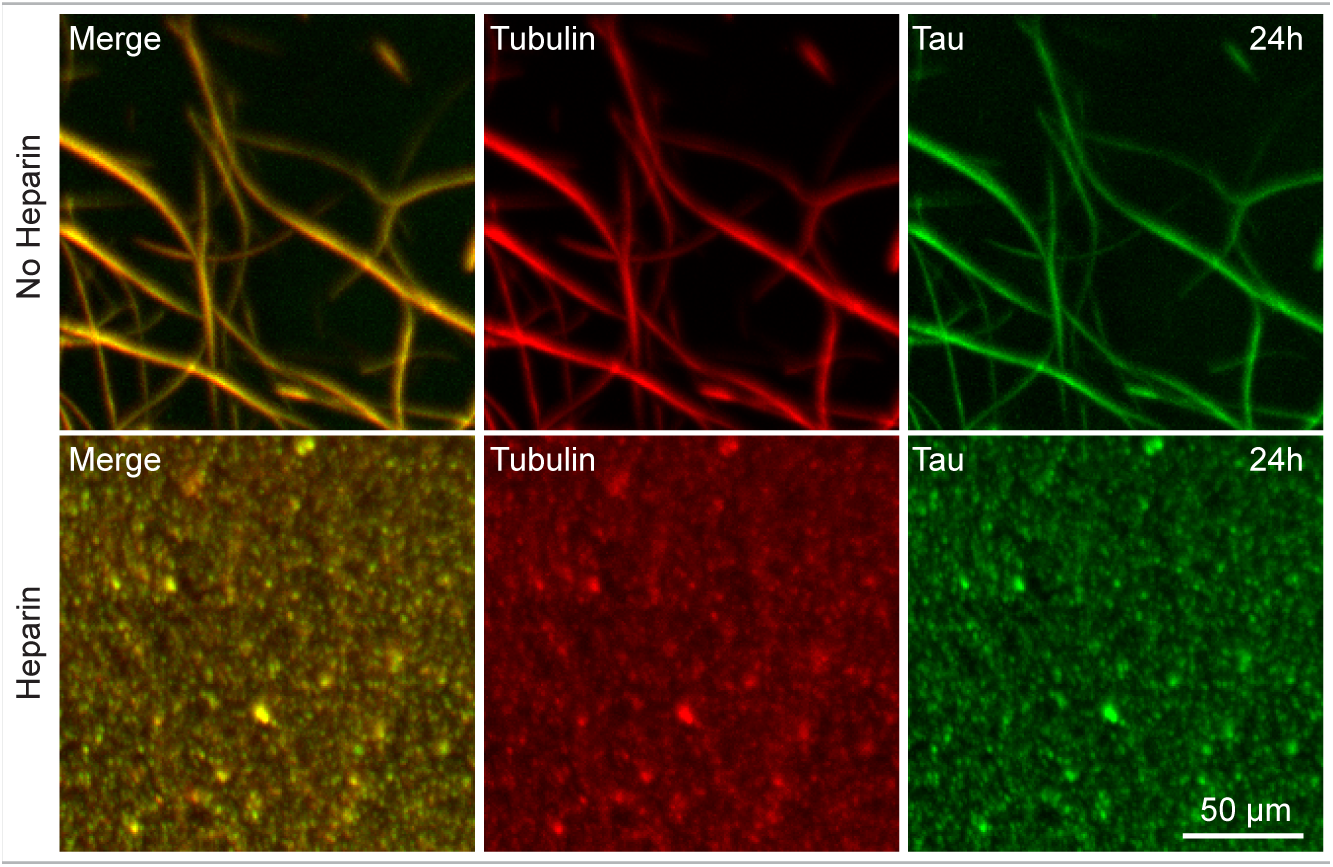
Tau-encapsulation stabilizes microtubules. Tau-encapsulated microtubule bundles with or without heparin observed 24 hours after formation. Bundles were formed from tau drops by addition of 5 μM rhodamine-tubulin and 1 mM GTP. Drops were formed with 25 μΜ of tau-EGFP in 1x BRB80 with 10% dextran and 1 mM DTT. Heparin was added at 200 μg/ml (final concentration) 30 minutes after bundles formation as indicated. Samples were incubated at room temperature for the duration of the experiment. Maximum projection confocal images are shown.

## SUPPLEMENTAL MOVIE LEGENDS

### Movie 1 - Fusion of tau drops using optical tweezer

Controlled fusion of tau drops using dual-trap optical tweezers. Two examples of the fusions events used for the analysis in figure 1C are shown. For each case, two droplets were trapped in two optical traps. While one trap was held stationary, the other trap was approached slowly until droplets were in close proximity, and fused shortly after. Smaller drops fused faster than bigger drops see inlet in figure 1C. Tau drops were formed with 25 μΜ tau-EGFP, 25 mM Hepes, 150 mM KCl, 1 mM DTT, 10% dextran, pH 7.4.

### Movie 2 - Internal rearrangement of tau drops

Time-lapse of the Fluorescent Recovery After Photo-bleaching (FRAP) after internal photo-bleaching of tau drops. Tau drops formed with 25 μM tau-EGFP, 25 mM Hepes, 150 mM KCl, 1 mM DTT, 10% dextran, pH 7.4. LUT, FIJI 16 colors. Scale bar, 2 μm.

### Movie 3 - Fission and surface wetting of tau drops

Tau drops fission and surface wetting upon an applied shear flow. Tau drops formed with 20 μM tau-EGFP, 25 mM Hepes, 150 mM KCl, 1 mM DTT, pH 7.4 with 10% dextran.

### Movie 4 - Tau-encapsulated bundles formation

Addition of 5 μΜ rhodamine-labelled tubulin together with 1 mM of GTP to tau drops. All drops deformed into rod-like shape within minutes after tubulin addition. Tau drops formed with 25 μΜ tau-EGFP with 10% dextran in BRB80 with 1 mM DTT.

### Movie 5 - Single drops deformation by internal microtubule bundles nucleation and polymerization

Detail of movie 4. Addition of 5 μΜ rhodamine-labelled tubulin together with 1 mM GTP to tau drops. A bidirectional deformation of the drops is observed with the polymerization of internal microtubule bundles. The volume of the drop is redistributed along the bundled arms as they extend. Tau drops formed with 25 μΜ tau-EGFP with 10% dextran in BRB80. Scale bar, 5 μm.

### Movie 6 - Single microtubules formation with soluble tau (no drops)

Addition of 5 μΜ of rhodamine-labelled tubulin together with 1 mM of GTP to 25 μΜ tau-EGFP in BRB80 with 1 mM DTT (no dextran added). Time-lapse of tau-EGFP signal.

### Movie 7 - Internal rearrangement of tau and tubulin in bundles

Time course of recovery of intensity (FRAP) after internal photo-bleaching of tubulin and tau in tau-encapsulated bundles. Bundles were formed by the addition of 5 μΜ of rhodamine-labelled tubulin together with 1 mM of GTP to tau drops. Tau drops formed with 25 μΜ tau-EGFP with 10% dextran in BRB80 with 1 mM DTT.

### Movie 8 - Fusion of Tau bundles

Fusion of three tau-encapsulated bundles. Bundles were formed by the addition of 5 μΜ of rhodamine-labelled tubulin together with 1 mM of GTP to tau drops. Tau drops formed with 25 μΜ tau-EGFP with 10% dextran in BRB80. Maximum projection fluorescent images of tau-EGFP are shown.

### Movie 9 - Tau coating zips single microtubules

Time-lapse of the zippering of two stable microtubules coated with tau-EGFP. 25 μΜ of tau-EGFP, 10% dextran in BRB80/Hepes buffer were flushed into a flow channel containing immobilized GMP-CPP stabilized microtubules. Scale bar, 10 μm.

### Movie 10 - Guided growth driven by preexisting microtubules

Guided growth of tau-encapsulated microtubules along preexisting microtubules. Tau drops were formed with 25 μΜ tau-EGFP and 10% dextran in BRB80. Cy5-tubulin (4 μΜ) together with 1 mM of GTP was added to the drops and flushed into a flow channel containing immobilized double stabilize rhodamine-labeled microtubules. Stabilized microtubules were attached to the surface by tubulin antibodies. Microtubules grow from drops (cyan) in the direction of the attached stable microtubules (magenta).

### Movie 11 - Tau encapsulation maintains microtubules bundled

Heparin addition to tau-encapsulated bundles interferes with tau-tubulin interaction. Time-lapse of the detachment of tau from microtubule bundles, its reshaping back into drops and the simultaneous debundling of microtubules after addition of heparin. Tau-EGFP shown in green and tubulin-rhodamine in red. 200 μg/ml of heparin was added to bundles formed from tau drops (25 μΜ tau-EGFP, 1x BRB80, 1 mM DTT, 10% dextran, pH 6.9, with 5 μΜ rhodamine-tubulin and 1 mM GTP added to drops). Maximum projection images.

